# Differing strategies used by motor neurons and glia to achieve robust development of an adult neuropil in Drosophila

**DOI:** 10.1101/149229

**Authors:** Jonathan Enriquez, Laura Quintana Rio, Richard Blazeski, Carol Mason, Richard S. Mann

## Abstract

In both vertebrates and invertebrates, neurons and glia are generated in a stereotyped order from dedicated progenitors called neural stem cells, but the purpose of invariant lineages is not understood. Here we show that three of the stem cells that produce leg motor neurons in *Drosophila* also generate a specialized subset of glia, the neuropil glia, which wrap and send processes into the neuropil where motor neuron dendrites arborize. The development of the neuropil glia and leg motor neurons is highly coordinated. However, although individual motor neurons have a stereotyped birth order and transcription factor code, both the number and individual morphologies of the glia born from these lineages are highly plastic, even though the final structure they contribute to is highly stereotyped. We suggest that the shared lineages of these two cell types facilitates the assembly of complex neural circuits, and that the two different birth order strategies – hardwired for motor neurons and flexible for glia – are important for robust nervous system development and homeostasis.

## Introduction

Cell lineages play a critical role in generating cellular diversity during animal development yet range widely in the number and types of cells they produce. Embryonic stem cells, for example, have the potential to give rise to all cell types, whereas hematopoietic stem cells have much more limited potential, giving rise only to myeloid and lymphoid cells (Seita and Weissman, 2010; Wu et al., 2016). The wide range of developmental potentials is also reflected in the degree to which cell lineages are hardwired or plastic in the types and numbers of cells they can generate. At one extreme, each of the 959 somatic cells of the adult *C. elegans* hermaphrodite is derived from an invariant lineage that can be traced back to the single-celled zygote (Sulston, 1976). In contrast, cell lineages of the mammalian immune system are plastic, and can be modified depending on the environmental challenges that an animal confronts (Boettcher and Manz, 2017). Similarly, intestinal stem cell lineages exhibit plasticity in response to injury and cell death (Apidianakis and Rahme, 2011; Jiang and Edgar, 2011).

Although the capacity of some cell lineages to respond to varying environmental conditions makes sense in light of the cell types these lineages generate, the underlying logic of why some lineages are invariant is usually not understood. For example, there is almost no correlation between lineage and cell type identity in *C. elegans*, where most sub-lineages contribute broadly to endodermal, mesodermal, and nervous system tissues. Consequently, a single cell type such as a motor neuron can be born from many different sub-lineages in *C. elegans* (Hobert, 2016). In *D. melanogaster*, although there are neuroblast (NB) stem cells that are dedicated to the generation of neurons, during embryogenesis each NB can generate a mixture of motor neurons, interneurons and glia that do not share an obvious function (Bossing et al., 1996; Schmidt et al., 1997). These observations raise the question of whether there is a biological reason for the structure of invariant lineages that may be difficult to discern by only examining the final progeny. Consistent with this notion, in the adult, the neuronal progeny of individual *Drosophila* NB hemilineages share anatomical features such as stereotyped axon paths and, when activated, the progeny of individual hemilineages can evoke specific behaviors in decapitated flies (Harris et al., 2015; Truman et al., 2004). Thus, by both anatomical and behavioral criteria, there is emerging evidence that individual *Drosophila* NBs generate functionally related progeny in the adult fly.

Another reason for the existence of hardwired lineages might be to facilitate the assembly of complex structures during development, which is an especially challenging problem for nervous systems. For example, the ~50 motor neurons (MNs) that innervate each leg in adult Drosophila are born from invariant NB lineages and have highly stereotyped birthdates and morphologies (Baek and Mann, 2009; Brierley et al., 2012). These stereotyped morphologies can be seen both in the muscles these MNs target and in their elaborate dendritic arbors that establish a very large number of neural synapses in dense neuropils present in each thoracic neuromere of the ventral nervous system (VNS) (Court, et al., 2017). Although each MN morphology is thought to be determined by unique combinations of morphology transcription factors (mTFs; (Enriquez et al., 2015)), how these stereotyped morphologies form the correct synaptic connections with interneurons and sensory neurons is not understood. Further, it remains an open question if the birth order of invariant NB lineages may also contribute to the assembly of complex neural circuits. Interestingly, at least one of the major NB lineages that give rise to adult MNs in the fly also generates another important cell type in the nervous system, glia (Baek et al., 2013; Lacin and Truman, 2016), which are critical for the establishment of neural synapses and the maintenance of neuronal activity (Freeman, 2015; Freeman and Rowitch, 2013). The observation that glia and leg MNs may be derived from common progenitors (neuroglioblast; NGB) raised the possibility that their development may be coordinated, and that being born from the same lineages might also play a role in neural circuit assembly. However, in contrast to MNs, very little is known about the glial types produced by these NGBs and how their morphologies are genetically specified.

In other contexts, *Drosophila* glia have been subdivided into three classes based on their functions and morphologies: surface glia, which isolate the CNS from the hemolymph, cortex glia, which send processes around neuronal cell bodies, and neuropil glia (NG), which surround neuropil and glomeruli (Omoto et al., 2016). NG are further subdivided into astrocytes and ensheathing glia (EG) (Awasaki et al., 2008; Muthukumar et al., 2014; Omoto et al., 2015; Peco et al., 2016). EG extend flat processes that surround neuropil and glomeruli as well as a small portion of axon bundles as they enter or leave the neuropil (Omoto et al., 2015; Peco et al., 2016). More recently, EG have been shown to extend small protrusions inside the neuropil of the adult brain (Kremer et al., 2017). In the medulla (region of the brain that processes visual information) EG are associated with axons (Kremer et al., 2017) and trachea (Kremer et al., 2017; Pereanu et al., 2007). In contrast, astrocytes send highly ramified processes deep into neuropil regions in close contact with synapses (Awasaki et al., 2008; Muthukumar et al., 2014; Stork et al., 2014). Although NG exhibit complex and diverse morphologies and are generally derived from stem cells that also generate neurons (Awasaki et al., 2008; Omoto et al., 2016), the developmental mechanisms leading to glial diversity remains unknown.

Here we show that three of the ~twelve lineages that generate leg MNs in *Drosophila* also generate all of the NG that populate the adult thoracic neuropil. As a consequence of being born from the same lineages, the development of the NG is physically and temporally coordinated with MN development. However, unlike the hardwired birth order and unique mTF codes of the MNs that are born from these NGBs, individual NG do not have a stereotyped morphology, birth order, or unique transcription factor code. Moreover, the gliogenesis phase of these lineages is plastic and highly adaptable: when gliogenesis in one lineage is compromised, other lineages compensate to maintain the correct number of NG. Thus, even though NG and MNs come from the same stem cells, and generate adult neuromeres with highly stereotyped structures, there are fundamental differences in how these two cell types are specified. We suggest that the combination of hardwired MN generation and flexible glia production facilitates the robust construction and homeostasis of complex neural circuits.

## Results

### Cellular organization of the adult thoracic neuropil glia

In order to characterize the cellular organization of the NG in the adult VNS, we expressed fluorescent reporters under the control of two Gal4 drivers, *alrm-Gal4* (astrocyte specific (Doherty et al., 2009)) and *R56FO3-Gal4* (EG specific (Kremer et al., 2017; Peco et al., 2016; Pfeiffer et al., 2008)), which we confirmed are expressed in all adult astrocytes and EG, respectively **(Figures 1 and S1)**. We also used as a marker Distalless (Dll), which is expressed in all NG in the adult VNS **(Figure S1)**.

**Figure 1.**
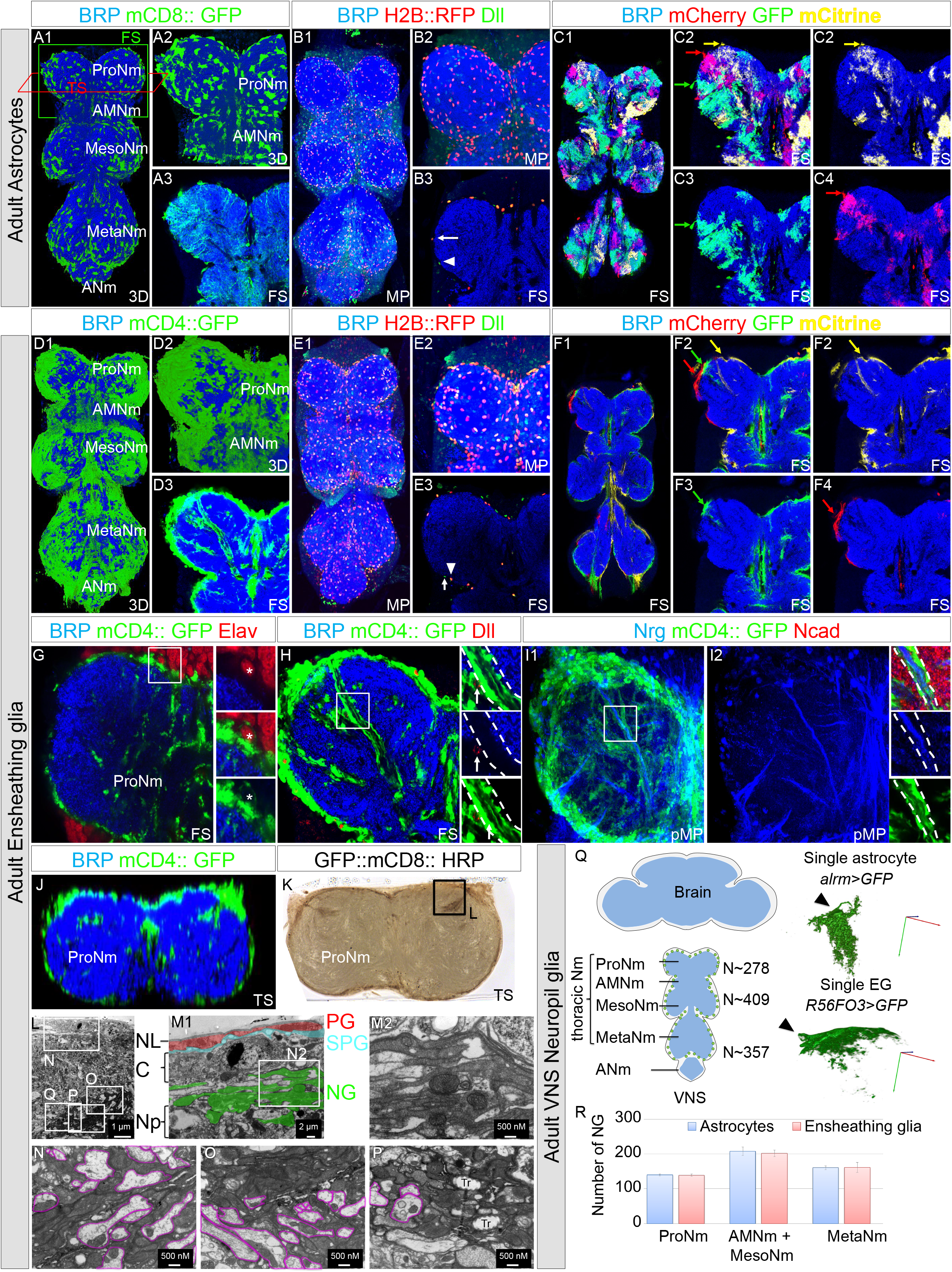
Cellular organization of the adult thoracic NG. **(A-C):** Adult VNSs immunostained with anti-BRP (neuropil marker, blue) **(A1-C4)** and anti-Dll (NG marker, red) **(B1-B3)** with astrocytes expressing mCD8::GFP (membrane marker, green) **(A1-C4),** H2B::RFP (Nuclear marker, red) **(B1-B3)** or the multicolor system FB1.1 (single cell marker, red, yellow and green) **(C1-C4)** under the control of *alrm-Gal4*. **(B1-B3)**, astrocyte (H2B::RFP+, Dll+): arrow, EG (Dll+): arrowhead. **(C1-C4)** Different astrocytes with processes occupying different neuropil territories: arrows. See also Figure S1 and video 1. **(D-F):** Adult VNSs immunostained with anti-BRP (neuropil marker, blue) **(D1-F4)** and anti-Dll (NG marker, red) (**E1-E3)** with EG expressing mCD4::GFP (membrane marker, green) **(D1-D3),** H2B::RFP (nuclear marker, red) **(E1-E3)** or the multicolor system FB1.1 (single cell marker, red, yellow and green; (Hadjieconomou et al., 2011)) **(F1-F4)** under the control of *R56F03-Gal4*. **(E1-E3),** astrocyte: arrow (Dll+), EG (H2B::RFP+, Dll+): arrowhead. **(F1-F4)** Different EG with processes occupying different neuropil territories: arrows. **See also Figure S1 and video 1.** **(G-I):** Prothoracic neuromeres with EG expressing mCD4::GFP and immunostained with anti-BRP (blue) and anti-Elav (neuron marker, red) **(G)**, anti-BRP (blue) and Dll (red) **(H)** or anti-NCad (neuropil marker, red) and Nrg (axon marker, blue) **(I1-2)**. **(G)** Asterisk: Elav+ neuron wrapped by an EG. **(H)** arrow: in more than half of the samples analyzed (N=10) a cell body of an EG (GFP +, Dll+) was observed inside the neuropil, next to axon bundles. Enlargements of the boxed regions are to the right of each panel. **See also video 2.** **(J-K):** Prothoracic neuromeres with EG expressing mCD4::GFP **(J)** and GFP::mCD8::HRP **(K)** immunostained with anti-BRP (blue) **(J)** or labeled with DAB (Black) **(K)**. **(L)**: Low magnification electron microscopy image of the boxed region in **(K)**. **(M-P):** enlargement of the boxed regions in **(L)**. PG: perineurial glia. SPG: subperineurial glia, NG: Neuropil Glia, C: Cortex, Np: Neuropil, NL: neural lamella. **(O)**: Left, schematic of adult CNS (blue: neuropils, grey: cortex); right, an individual astrocyte and EG labeled with mCD8::GFP under the control of *alrm-Gal4* and *R56F03-Gal4* using the MARCM technique. Axes: green (posterior), blue (dorsal), red (medial). **(R):** Average number of NG in the adult VNS, in which EG or astrocytes were expressing H2B::RFP under the control of *R56F03-Gal4* or *alrm-Gal4*, respectively, and immunostained with anti-Dll. Number of samples = 4/genotype. Error bars indicate standard deviation. ProNm: Prothoracic neuromere, AMesoNm: Accessory mesothoracic neuromere, MesoNm: Mesothoracic neuromere, MetaNm: Metathoracic neuromere, ANm: Abdominal neuromeres; FS: frontal cross section, TS: transverse cross section, 3D: 3 dimensional reconstruction of confocal image stack. pMP: partial maximum projection.

As for the NG in the larval CNS (Beckervordersandforth et al., 2008; Omoto et al., 2015), the number of NG surrounding each thoracic neuropil is very consistent between animals (with a very low standard deviation; **Figure 1Q, R).** However, there are ~30-fold and ~10-fold more NG in one adult thoracic neuropil than in a larval hemisegment or a larval brain hemisphere, respectively **(Figure 1Q, Supp. Table 1)**. These differences correlate with the dramatically different sizes and complexities of these structures. Underscoring this point, if we compare the number of adult NG in the adult thorax (~2100) with the number of NG in the adult brain (~6800; (Kremer et al., 2017)), nearly one fourth of all the NG in the adult CNS are localized in the thorax.

As described for other regions of the CNS (Awasaki et al., 2008; Muthukumar et al., 2014), adult thoracic astrocytes send processes deep into the neuropil that respect each other’s territory, thus exhibiting a tiling-like phenomenon **(Figure 1A3, C, Video 1).** EG extend processes that surround each neuropil and also respect each other’s territory with very infrequent overlaps **(Figure 1D3, F, Video 1)**. In addition, the adult thoracic EG feature a more complex morphology than previously described for larval EG. First, adult EG wrap the cell bodies of neurons (Elav+) that are localized next to the neuropil **(Figure 1G, Video 2)** as well as astrocyte cell bodies **(Figure S1)**. Second, they send processes inside the neuropil where they wrap axon bundles (Neuroglian+) where no synapses are formed (BRP-negative, a marker of the active presynaptic zone) **(Figure 1H, I, video 2)**. Interestingly, EG only wrap these axons inside the neuropil; outside of the neuropil these axons are surrounded by a distinct class of wrapping glia **(Figure 1H and data not shown)**. Thus, EG isolate nerves inside neuropil before they make synapses with other neurons.

We next performed electronic microscopy of transverse sections of prothoracic neuromeres (ProNs) **(Figure 1L-P).** EG wrap the thoracic neuropil very densely with several layers **(Figure 1M)**. These results, in combination with the tiling results described above, suggest that one EG can send processes organized in layers around the neuropil. Inside the neuropil, axon bundles and single axons can be wrapped by layers of EG processes **(Figure 1N-P)**. Interestingly, each nerve also contains trachea bundles that are surrounded by EG, forming a structure with all three cell types: EG, axons, and trachea **(Figure 1P)**. Notably, axons close to the midline are not wrapped by EG (data not shown) but are wrapped instead by another category of glia that may be a specialized form of trachea EG (Kremer et al., 2017).

## Developmental origins of the adult thoracic NG

As described above, the adult thoracic neuropils are complex structures containing glia processes and axons/dendrites in close juxtaposition to each other, raising the question of how these structures develop. A previous study suggested that pockets of glia, marked by the general glia marker Repo, surround the immature leg neuropil in late 3^rd^ instar larvae (just prior to the onset of metamorphosis) and produce adult astrocytes (Li et al., 2014), while another study suggests that these glia express Dll and may migrate to the leg to form wrapping glia, which are distinct from EG and astrocytes (Plavicki et al., 2016). In order to definitively determine what type of adult cells these Repo+ glia give rise to, we used a *Dll-Gal4* enhancer trap and a Dll antibody to label these cells during development **(Figure 2)**.

**Figure 2.**
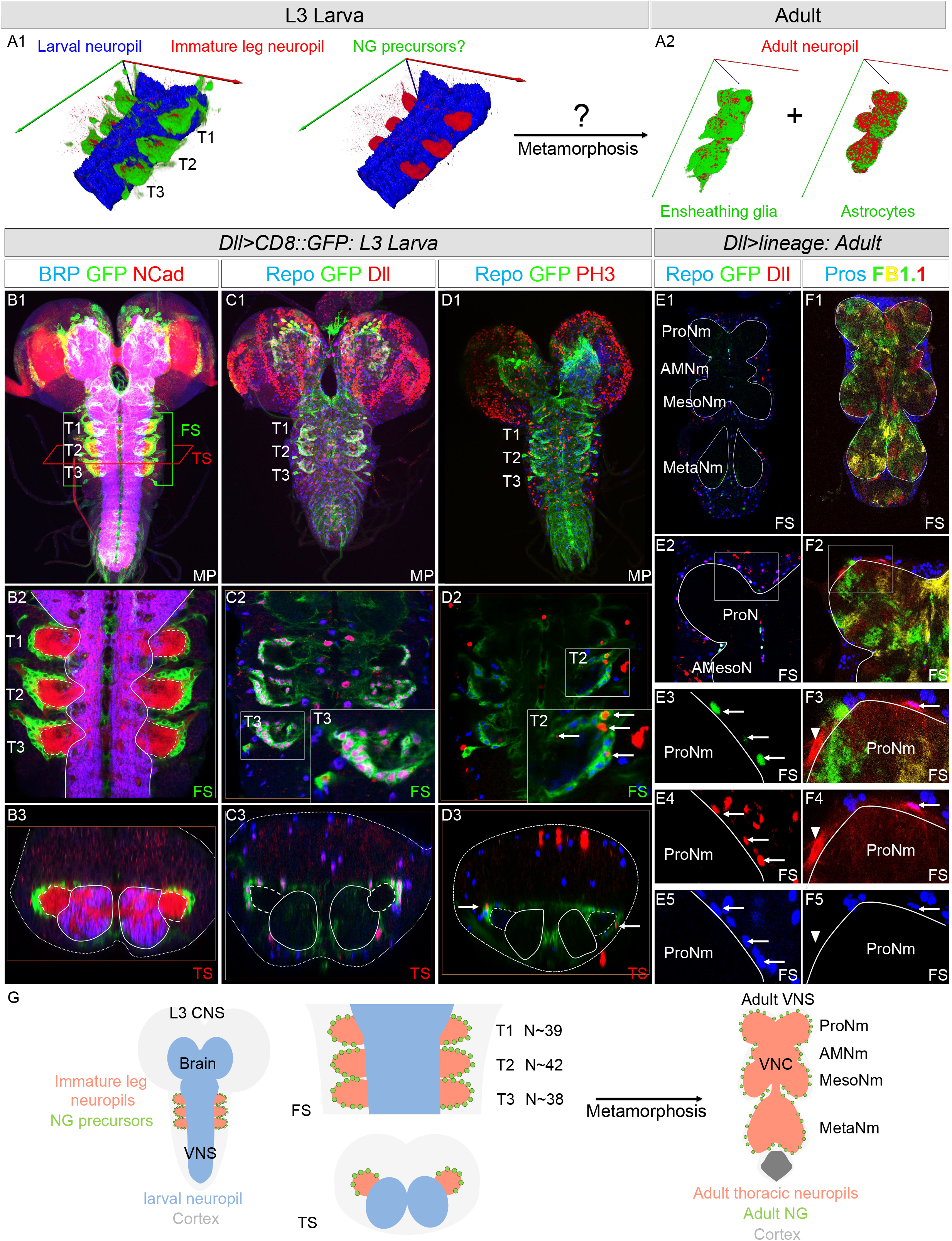
Developmental origin of the adult NG. **(A1)**: 3D rendering of the thoracic segments of a L3 VNS labeled with anti-BRP (mature neuropil marker, blue) and anti-NCad (mature and immature neuropil marker, red). The glia surrounding the six immature leg neuropil express mCD8::GFP (green, left image only) under the control of *Dll-Gal4*. (T1, T2, T3 indicate the three thoracic segments). **(A2)**: 3D rendering of adult VNSs with astrocytes and EG expressing GFP under the control of *alrm-Gal4* (right) and *R56F03-Gal4* (left) and costained with anti-BRP (red). **(B-D)**: L3 CNSs in which glia surrounding the six immature leg neuropil express mCD8::GFP under the control of *Dll-Gal4* and costained with anti-BRP (blue), anti-NCad (red) **(B1-B3)**, anti-Repo (blue), anti-Dll (red) **(C1-C3)**, or anti-Repo (blue), anti-PH3 (red) **(D1-D3)**. **(D2-D3)** arrows point to glia expressing PH3. **(E-F)**: Adult VNSs with neuropil glia expressing nuclear GFP (green) (E1-E5) or the multicolor Flybow system FB1.1 (yellow, red and green) (see **Methods** for details) and immunostained with anti-Repo (blue), anti-Dll (red) (**E1-E5)** or anti-Pros (Blue) (**F1-F5).** (**E1-E5),** arrow points to adult NG expressing GFP. **(F1-F5),** arrow: astrocyte (mCherry+, Pros+), arrowhead: EG (mCherry+, Pros-). See also **Figure S2**. **(G)**: Left: schematic of an L3 CNS. The immature leg neuropil and NG precursors are indicated. Right: schematic of an adult VNS with the adult leg neuropil glia. N, average number of NG precursors in each L3 thoracic hemisegment.

In 3^rd^ instar larvae (L3), there were ~40 Repo+ Dll+ cells surrounding each immature leg neuropil **(Figure 2B, C, G)**, where nascent MN axons enter and then exit

**(Figure 4A, I)**. If these cells are the progenitors of the adult NG they should be mitotically active. To test this idea, we first confirmed that they express phosphorylated histone H3 (pH3), a marker for mitotically active cells **(Figure 2D)**.

Although Dll protein is expressed in adult NG, *Dll-Gal4* is only active in immature NG **(Figure S4)**, allowing us to mark the progeny of *Dll-Gal4* expressing larval cells in lineage tracing experiments **(see Methods)**. In the first experiment, we labeled the adult progeny of *Dll-Gal4-* expressing L3 glia with nuclear GFP (*UAS-nGFP*) and co-stained with an antibody against Repo and Dll. **(Figure 2E)**. Only the adult Repo+ Dll+ NG were labeled with nGFP, demonstrating that the *Dll-Gal4* expressing cells in the larval CNS only generate adult NG. In a second experiment, we used *Dll-Gal4* to activate the Flybow system (*UAS-FB1.1*), which permanently labels cells with different fluorescent markers (Hadjieconomou et al., 2011). We co-stained the resulting adult VNSs with Prospero, which is expressed in astrocytes but not in EG (Griffiths and Hidalgo, 2004; Peco et al., 2016) **(Figure 2F and Figure S1)**. Based on Prospero expression and morphology, we found that both astrocytes and EG were labeled. Significantly, the Flybow system did not label other types of glia, such as surface glia or cortex glia, arguing that these glia subtypes are not derived from *Dll-Gal4-*expressing cells. We confirmed this result with another driver (*R31F10-Gal4*) that is also expressed in immature NG **(Figure S2)**.

Taken together, we conclude that the ~40 Repo+ Dll+ *Dll-Gal4-*expressing cells surrounding each immature leg neuropil in L3 are the precursors of the >280 NG that surround each adult thoracic neuropil.

### Coordinated development of the adult NG and neuropil

To characterize the development of the adult neuropil, we conducted several experiments to trace the origins of this complex structure **(Figure 3)**.

**Figure 3.**
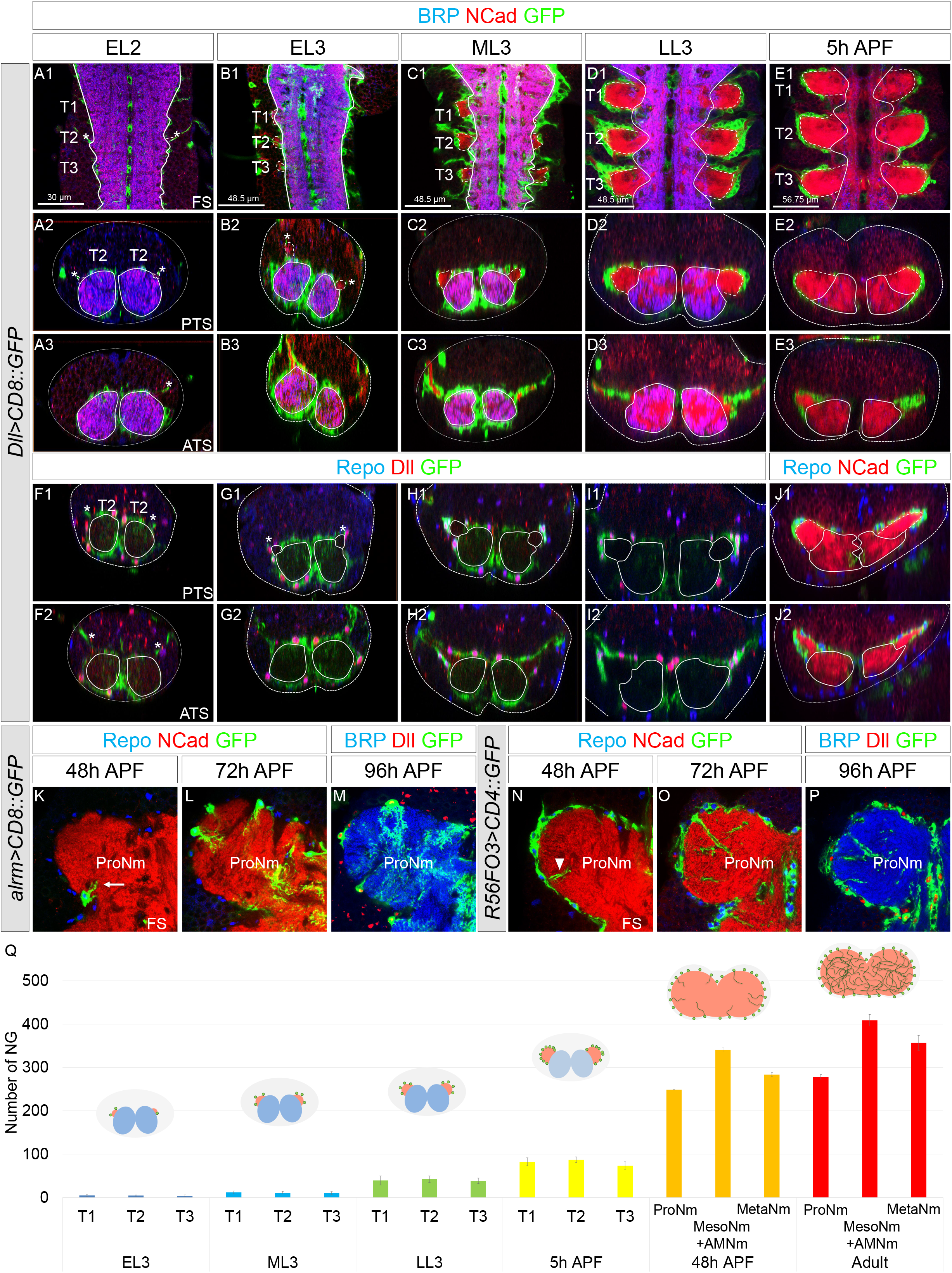
Coordinated development of NG and adult neuropil. **(A-J):** Thoracic segments of larval VNSs at different stages with NG precursors expressing mCD8::GFP and labeled with anti-BRP (blue), anti-NCad (red) **(A1-E3),** anti-Repo (blue), anti-Dll (red) **(F1-I2)**, or anti-Repo (blue), anti-NCad (red) **(J1-J2)**. Asterisks indicate axons of incoming leg sensory neurons. See also **Figure S3** and **video 3** and **4**. (**E1-E3**) At this stage, the expression of BRP is almost undetectable, probably a consequence of the remodeling and/or degradation of the larval neuropil. **(K-P):** ProNm of an adult VNS with astrocytes **(K-M)** or EG **(N-P)** expressing a membrane-tagged GFP under the control of *alrm-Gal4* and *R56F03-Gal4*, respectively, and costained with anti-Repo (blue) and anti-NCad (red) **(K,L,N,O)**, anti-BRP (blue) and anti-Dll (red) (**M,P**). **(K,N)** Processes of astrocytes (arrow) and EG (arrowhead) are first observed invading the neuropil. **(Q):** Average number of NG at different stages (number of samples analyzed at each stage = 4) with schematics of transverse sections of T1 or ProNm. NG (green), larval neuropil (blue) and leg neuropil (red) are indicated. Error bars indicate standard deviation. ATS: Anterior Transverse Section, PTS: posterior transverse section. APF: after pupa formation

In early 2^nd^ instar larvae (EL2), most of the adult leg MNs are not yet born (Baek and Mann, 2009) and no *Dll>GFP+* NG precursors are visible. However, at this stage incoming leg sensory axons are visible and mark the location where the adult neuropil will form **(Figure 3A, F)**. In early L3 (EL3; 72 hrs after egg laying, AEL), 0-10 NG precursors are observed per thoracic hemisegment **(Figure 3B, G and Q)**, and their numbers gradually expand through subsequent larval and pupal stages (**Figure 3C-E, H-Q)**. As NG precursors are produced they send processes that surround the growing neuropil and wrap the leg nerves **(Figure 3A-J)**. By the mid pupal stage (48hr APF), most of EG and astrocytes have been generated and start to differentiate by sending processes inside the neuropil **(Figure 3K-N)**. By the late pupal stage (96hr APF) the adult neuropils are fully invaded by astrocyte processes and wrapped by EG **(Figure 3M, P)**.

These results suggest that the adult thoracic NG are produced from early L3 to mid pupa. These glia create a structure that facilitates the maturation of the adult neuropil **(Video 4)**; by L3, the MN neurites are completely surrounded by the processes of the immature NG. During metamorphosis, the dendrites and axons of the neurons establish their final morphologies within the glia-defined neuropil.

### The adult thoracic NG are produced by 3 neuroglioblasts that also generate leg motor neurons

Previous work established that the larval NG are born from a single glioblast in each hemisegment of the larval CNS (Awasaki et al., 2008; Beckervordersandforth et al., 2008; Ito et al., 1995; Jacobs et al., 1989) and from a single glioblast or neuroglioblast (NGB) in each hemisphere of the brain (Hartenstein et al., 1998; Omoto et al., 2015). To characterize the lineages that give rise to the adult thoracic NG, we carried out several clonal analysis experiments. Previous observations demonstrated that NB lineages such as Lin A (also called Lin 15), produce both leg motor neurons and glial cells (Baek et al., 2013; Truman et al., 2004). This observation raised the intriguing possibility that NG and MNs may share common developmental origins.

First, we used MARCM to express the membrane reporter *UASmCD8::GFP* with two different drivers, *repo-Gal4* (expressed in all glia, (Sepp and Auld, 2003)) and *DVGLUT-Gal4* (expressed in all MNs and some interneurons (Mahr and Aberle, 2006)). We induced these clones in L1 larvae and dissected late L3 CNS and co-stained for Dll, to mark immature NG, and BRP to mark the mature neuropil. If a stem cell produces both MNs and NG, individual MARCM clones should contain both cell types. We identified glial cells by their position and expression of Dll, and MNs by their morphology, particularly MN axons, which exit the CNS and target the leg imaginal discs (Baek and Mann, 2009; Truman et al., 2004)). 62/62 clones that labeled NG also labeled MNs **(Figure 4)**. We found 3 types of clones based on the number of MNs they contained. Type 1 clones (N=36) contained ~29 leg MNs and ~20 immature adult NG. Based on MN morphology, we concluded that this lineage is Lin A **(Figure 4A)**. Type 2 clones (N=23) contained 2 MNs and ~10 immature NG **(Figure 4B)**. In type 3 (N=3), the clones contained 1 MN and ~7 glia **(Figure 4A)**. The total number of glia (~37) produced by these three NGBs is very close to the total number of NG progenitors that surround each neuropil at L3 (~40), consistent with the idea that these three lineages produce most, and perhaps all, of the adult NG.

**Figure 4.**
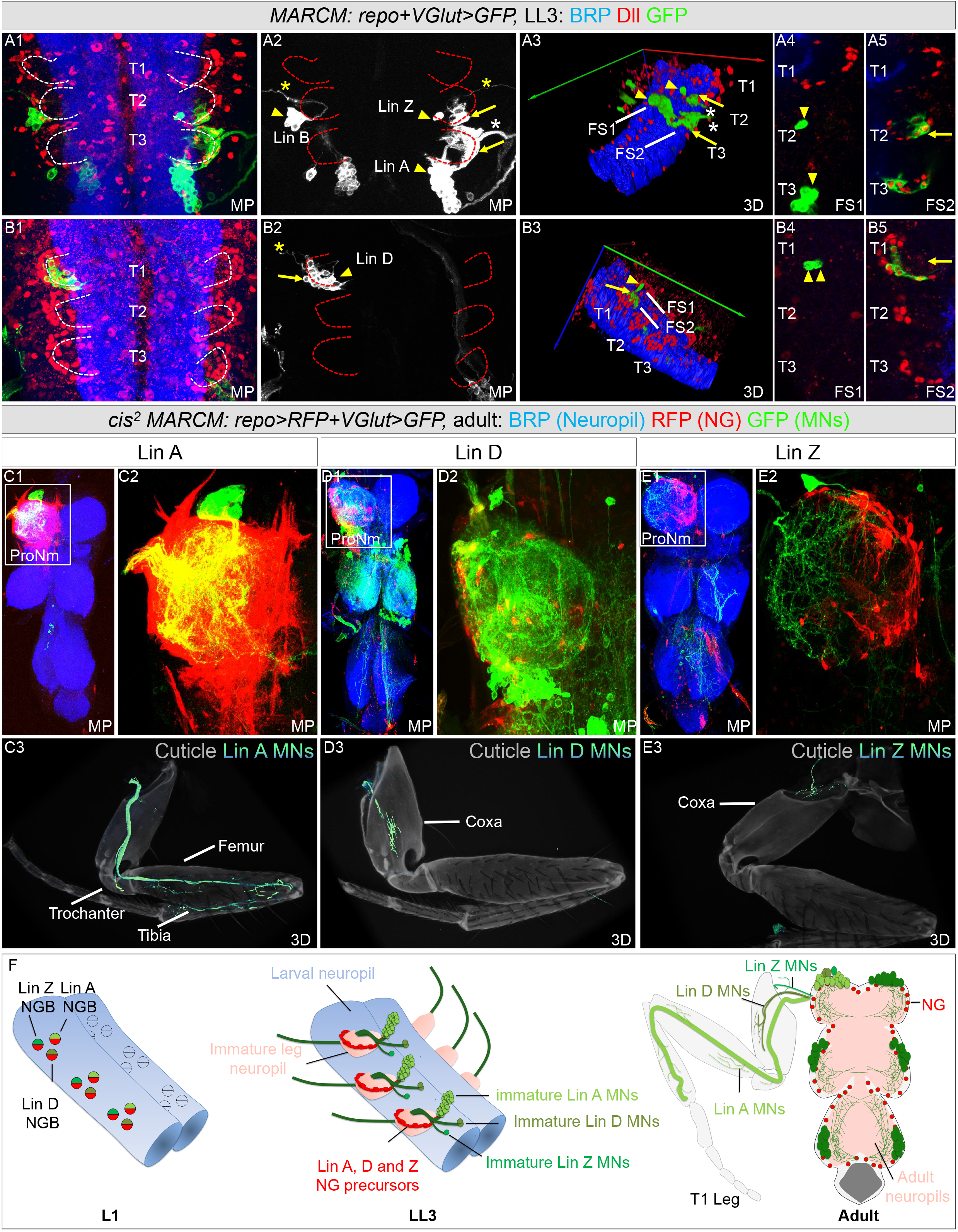
3 NGBs per thoracic hemisegement produce leg MNs and all NG. **(A-B)** Thoracic segments of an L3 VNS labeled with anti-BRP (blue) and anti-Dll (red) containing MARCM clones expressing mCD8::GFP under the control of both *VGlut-Gal4* and *repo-Gal4.* Lin A, Lin Z, and Lin D clones produce MNs (arrowhead) and glia (arrow). **(A1-A5)** Sample with Lin A, Lin B and Lin Z clones; note that Lin B does not produce glia. **(B1-B5)** Sample with a Lin D clone. Asterisks mark MN axons exiting the VNS. **(C-E):** Lin A **(C1-C3)**, Lin D **(D1-D3)** and Lin Z **(E1-E3)** *cis^2^* MARCM clones in a ProN labeled with anti-BRP (blue) **(C1-C2, D1-D2, E1-E2)** and in a T1 Leg **(C3, D3, E3)** expressing mCD8::RFP in NG (red) and mCD8::GFP (green) under the control of *repo-Gal4* and *VGlut-QF*, respectively. **(F):** Left: schematic of the thoracic segments of an L1 VNS with the Lin A, Lin B, Lin Z NGBs; Center: schematic of the thoracic segments of a late L3 VNS with Lin A, Lin B, Lin Z clones; Right: schematic of an adult VNS with Lin A, Lin B, Lin Z where MNs and NG are labeled in green and in red respectively. Note that only the innervation of a T1 leg is shown.

In a second approach, we generated clones using a modified QMARCM/MARCM method (*cis^2^-*MARCM) in which the two transcriptional repressors, GAL80 and QS, are recombined on the same chromosome arm (see **Methods** for details). We used two drivers, *DVGLUT-QF* and *repo-Gal4*, allowing us to visualize VGlut+ MNs and Repo+ glia derived from the same stem cell with two different fluorescent markers. Using this method, and analyzing the adult leg and VNS, we confirmed that there are three NGBs that produce leg MNs and adult NG. Lin A clones produce NG and ~29 MNs targeting the trochanter, femur and tibia (N=14, in ProN) **(Figure 4C, F)**. Type 2 clones labeled NG and 2 MNs that targeted the coxa; based on the morphology of these MNs, we recognized this lineage as Lin D (Baek and Mann, 2009) (N=2) **(Figure 4D, F)**. Type 3 clones labeled NG and a single MN that targeted a body wall muscle at the base of the coxa (N=16) **(Figure 4E, F)**. To our knowledge, this lineage has not been characterized previously so we named it Lin Z.

Third, to determine if these three lineages generate *only* MNs and NG, we repeated the MARCM experiments using a tubulin driver (*tub-Gal4*), which labels all cell types. We stained the resulting L3 larvae with Elav (a pan neuronal marker), Deadpan (a NB marker) and Repo (a glia marker) (**Figure 5)**. These results confirmed that Lin A generates MNs and NG. Lin A clones usually included two cells that did not stain for either Elav or Repo. Because of their proximity to the NB, they are likely to be ganglion mother cells (GMCs) **(Figure 5A and data not shown)**. All non-Lin A clones (N=4) that generated Repo+ NG labeled only 1 or 2 Elav+ neurons **(Figure 5 B, C).** Together with our earlier clonal experiments, these results suggest that, post-embryogenesis, Lin D, Lin Z, and Lin A give rise to only leg MNs and NG. Notably, a Deadpan+ NB was always associated with Lin A clones, but not with Lin D or Lin Z clones

**Figure 5.**
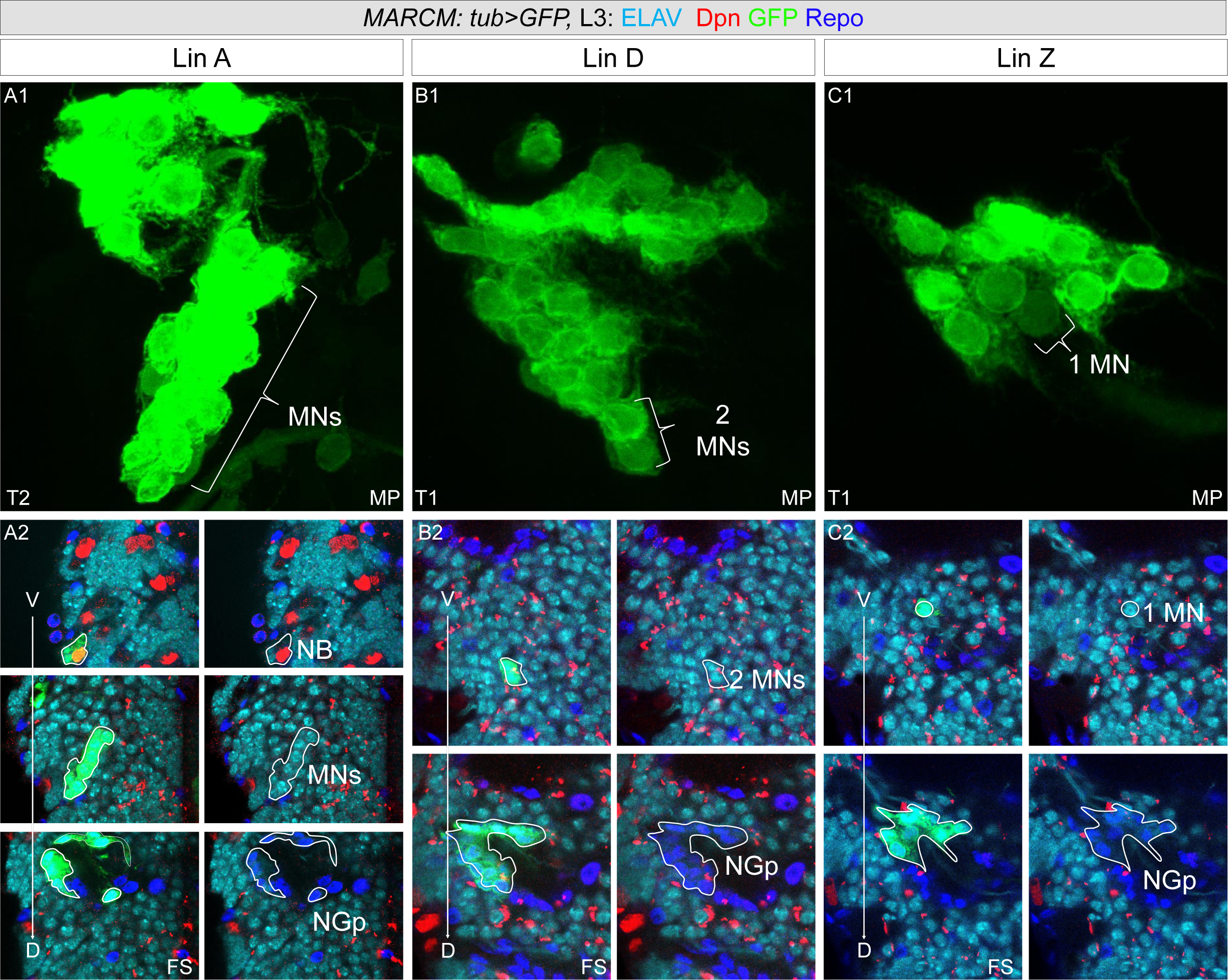
The 3 NGBs do not produce cell types other than MNs and NG. **(A-C):** Lin A **(A1-A2)**, Lin D **(B1-B2)**, Lin Z **(C1-C2)** MARCM clones labeled with mCD8::GFP and nGFP under the control of *tub-Gal4* in a thoracic segment of an L3 VNS and immunostained with anti-Dpn (neuroblast marker, red), anti-Elav (cyan) and anti-Repo (blue). **(A2, B2, C2)** Frontal Sections (FS) from ventral (V) to dorsal (D). NGp: neuropil glia progenitor (Repo+)

Finally, to determine which types of NG are produced by these three lineages, we repeated the *cis^2^*-MARCM experiments with *alrm-Gal4* or *R56FO3-Gal4* (instead of *repo-Gal4*), to label astrocytes and EG, respectively **(Supp. Table 2)**. We found that Lin A (N=14) and Lin Z (N=19) produce both types of NG while Lin D (N=4) only generates astrocytes **(Figure 6, Supp. Table 2).**

**Figure 6.**
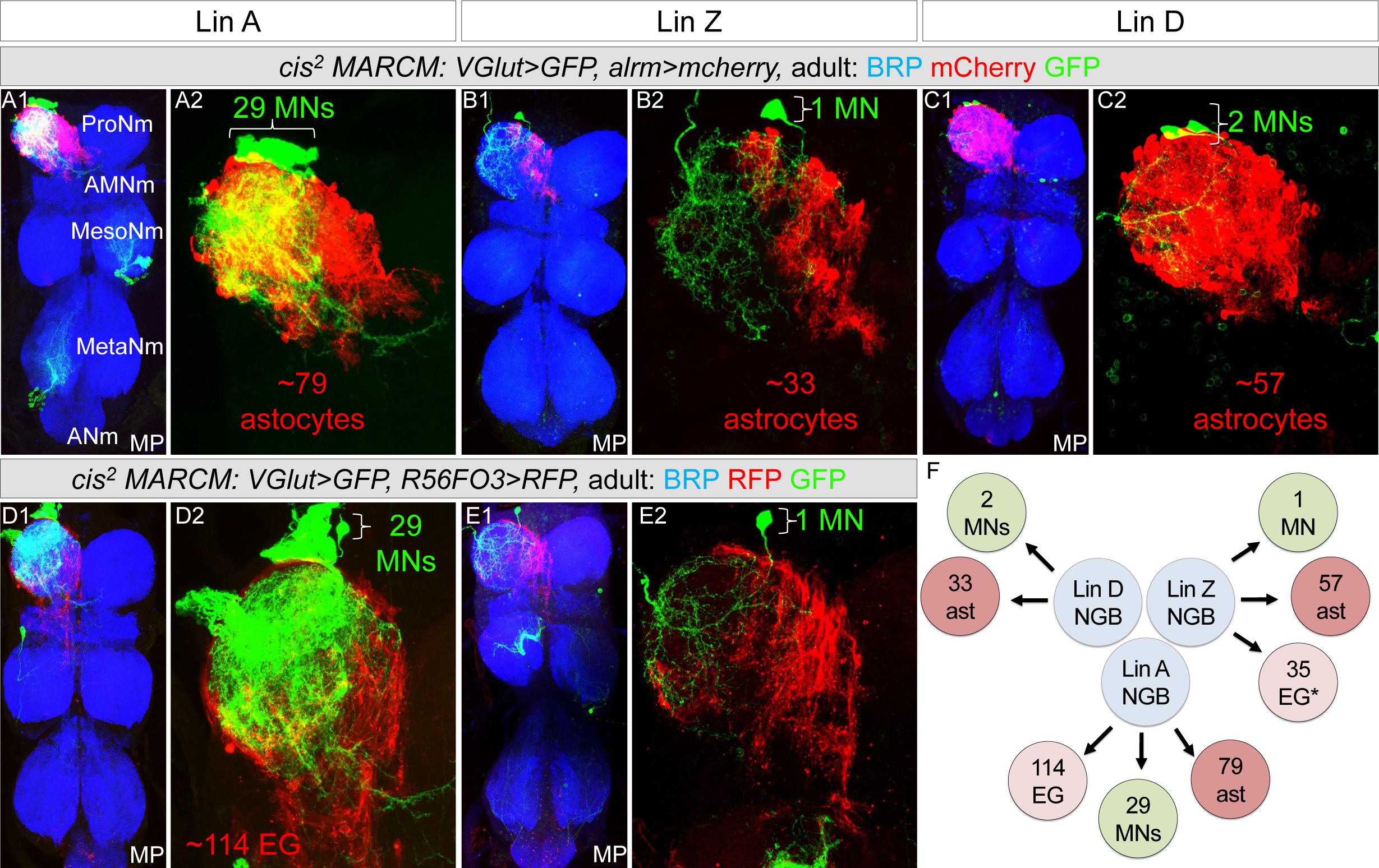
Lin A and Lin Z produce astrocytes and NG while Lin D produces only astrocytes. **(A-E):** Lin A (**A1-A2, D1-D2**), Lin Z (**B1-B2, E1-E2**) and Lin D (**C1-C2**) *cis^2^* MARCM clones in a ProNm labeled with anti-BRP (blue) with MNs expressing mCD8::GFP (**A1-E2**), astrocytes expressing mCherry (**A1-C2**) and EG expressing mCD8::RFP (**D1-E2**), under the control of *VGlut QF*, and *alrm-Gal4* or *R56F03-Gal4*, respectively. **(F):** Summary of progeny produced by Lin A, Lin D, and Lin Z. The numbers are averages for each NGB in a ProNm. Number of VNSs analyzed: Lin A astrocyte (ast): N=10, Lin A EG: N=4, Lin D astrocyte: N=4, Lin Z astrocytes: N=13. Note: the number of EG produced by Lin Z (*) is an estimate obtained by subtracting the number of EG produced by Lin A from the total number of EG.

We conclude that in each thoracic hemisegment, all of the adult NG are generated from three NGBs, which also give rise only to leg MNs.

### Expression and function of Dll during NG development

In the thoracic neuromeres, Dll is expressed in one pair of interneurons per hemisegment and at different levels in subpopulations of glia: Dll is expressed at high levels in EG and astrocytes, in low levels in cortex glia, and is not expressed in surface glia **(Figure S1, S4)**. These differences in Dll levels are also observed in the thoracic segments of L3 larvae. In addition, Dll is expressed at very high levels in the immature adult NG glia at the L3 stage **(Figure S4)**.

To assess if Dll plays a role in NG development, we analyzed the morphology and number of NG produced by *Dll* mutant Lin A clones. We found that, in the adult VNS, the number of both types of NG is reduced in *Dll* mutant clones, but the morphology of the surviving NG appears normal **(Figure 7, S4)**. Interestingly, in L3 larvae the number of immature NG in *Dll* mutant Lin A clones (~20) is normal **(Figure S4)**, suggesting that *Dll* mutant NG divide more slowly and/or are eliminated during metamorphosis. The morphology (dendritic arbor and axonal targeting) and number of MNs produced by Lin A are not affected in *Dll* mutant clones, consistent with the lack of *Dll* expression in Lin A MNs.

**Figure 7.**
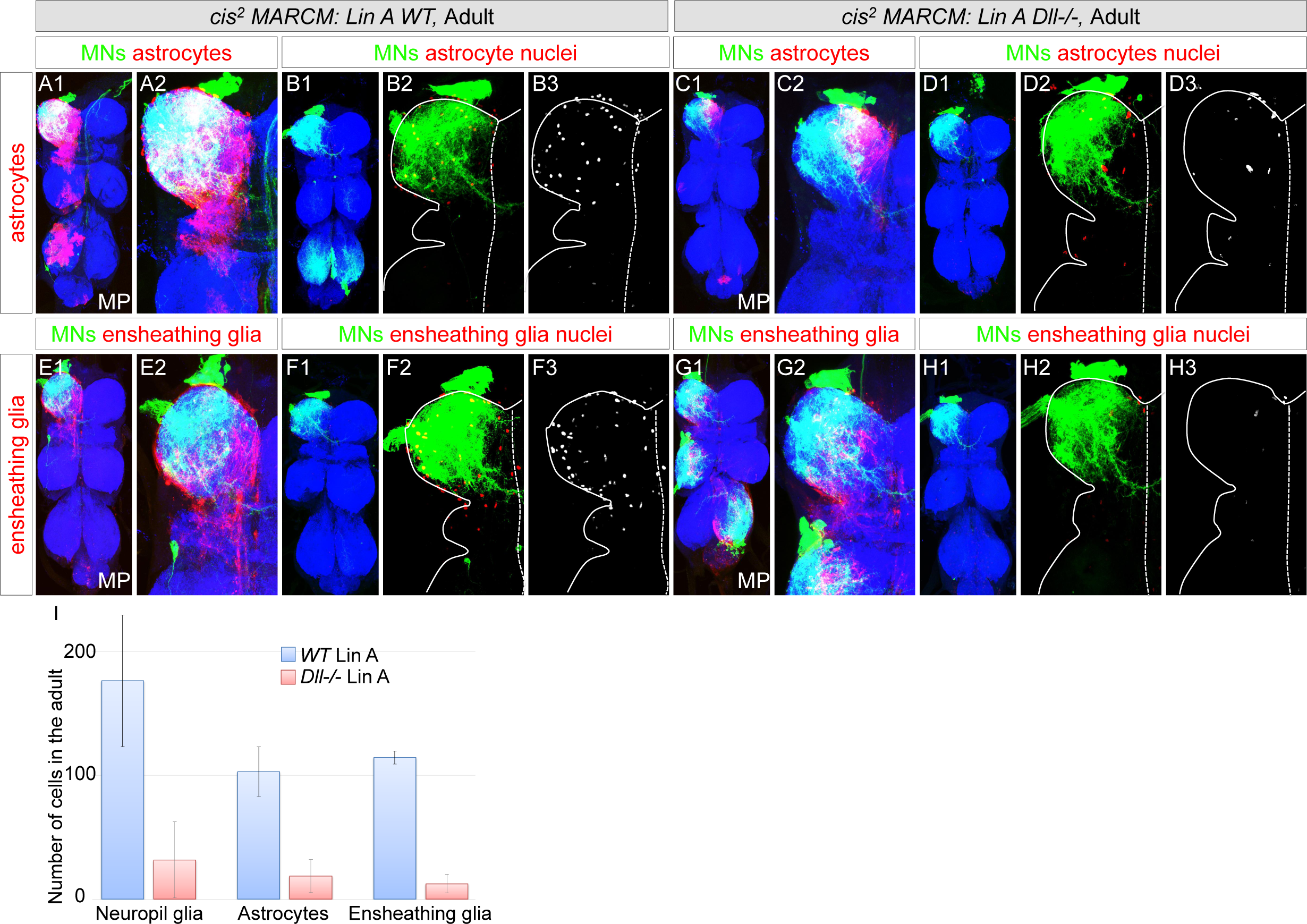
Expression and function of Dll during NG development. **(A-H)** *WT* **(A1-B3, E1-F3)** and *Dll* **(C1-D3, G1-H3)** *cis^2^* MARCM Lin A clones expressing mCD8::GFP **(A1-H3)**, mCherry **(A1-A2, C1-C2, E1-E2, G1-G2)**, H2B::RFP **(B1-B3, D1-D3, F1-F3, H1-H3)** under the control of *alrm-Gal4* **(A1-D3)** and *R56F03-Gal4* **(E1-H3)** and costained with anti-BRP (blue). **(I)** Average number of *repo>H2B:RFP+* neuropil glia, *alrm>H2B:RFP+* astrocytes and *R56F03>H2B:RFP+* EG in *WT* (blue bar) and *Dll-/-* (red bar) *cis^2^* MARCM Lin A clones. Error bars indicate standard deviation. See also Figure S5.

### Unlike MNs, individual NG born from the same lineage are not stereotyped

Previous studies have revealed that each of the 47 leg MNs have a specific birth date and morphology, characterized by their axonal targeting, dendritic arbor and cell body position (Baek and Mann, 2009; Brierley et al., 2012). We recently characterized Lin B, a lineage that produces only 7 MNs, and described a post-mitotic code of mTFs that controls individual MN morphologies (Enriquez et al., 2015). We identified these TFs by an expression screen using ~250 antibodies yet, surprisingly, none of these TFs are differentially expressed in subsets of thoracic NG (data not shown). Instead, we found TFs, such as Dll, expressed in all NG. The one exception is Prospero (Pros), which is expressed in all astrocytes but not EG (Kato et al., 2011; Griffiths and Hidalgo, 2004; Peco et al., 2016) **(Figure S1)**. If no single cell TF code exists for NG, we reasoned that the number and morphology of the NG progeny from the same lineage may appear different from animal to animal. Below we used the *cis^2^*-MARCM technique to test this prediction **(Figure 8A-R, Supp. Table 3)**.

**Figure 8.**
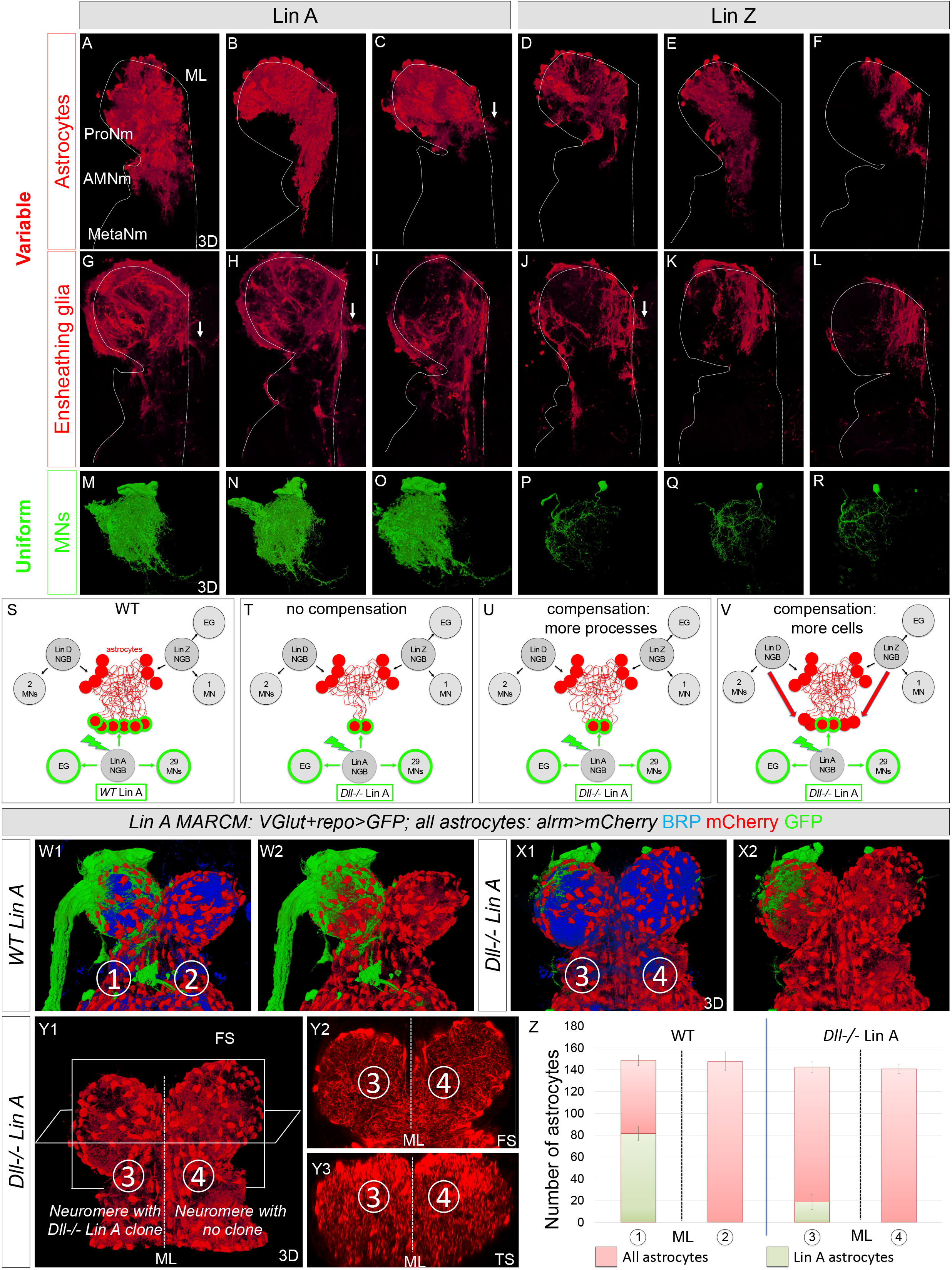
Gliogenesis is not stereotyped and depends on competition. **(A-R)** Lin A **(A-C, G-I, M-O)** and Lin Z **(D-F, J-L, P-R)** *cis^2^* MARCM clones expressing mCherry **(A-F)** or mCD8:RFP **(G-L)** and mCD8::GFP **(M-R**) under the control of *alrm-Gal4* **(A-F)**, *R56F03-Gal4* **(G-L)** and *VGlut-QF* **(M-R)**. Note, all clones **(A-L)** express mCD8::GFP (not shown). **(M-R)** show three examples each of the MN progeny for Lin A (**M-O**) and Lin Z (**P-R**). Astrocyte and EG clones are variable from animal to animal, while MN clones have a uniform, lineage-specific morphology. **(C)** Arrow points to astrocyte processes crossing the midline. **(G,H)** Arrows point to EG processes crossing the midline. **(H)** Arrow points a EG cell body on the contralateral neuromere. **(S)** Schematic summarizing the Lin A, Lin D and Lin Z NGBs and their progeny in a thoracic neuromere. Green indicates a Lin A MARCM clone induced in the NGB (lightning bolt). **(T-V)** Same as **(S)** showing three possible outcomes for a *Dll-/-* Lin A MARCM clone: no compensation **(T)**, an increase in astrocyte processes **(U)** or an increase in astrocyte number **(V)**. **(W-Y)** Thoracic neuromeres with WT (W1-W2) and *Dll-/-* (X1-Y2) Lin A MARCM clones in the right ProNm, labeled with mCD8::GFP (green) under the control of *VGlut-Gal4* and *repo-Gal4*, and costained with anti-BRP (blue). All astrocytes are labeled with mCherry under the control *alrm QF*. **(Y1-Y3)** Frontal (FS) or transverse (TS) sections showing that all thoracic neuropil appear to be equally invaded by astrocyte processes even if the neuropil contains a *Dll-/-* Lin A clone. **(Z)** Average number of astrocytes in *WT* and *Dll-/-* MARCM Lin A clones (GFP+, mCherry+), compared to the number of astrocytes produced by Lin D and Lin Z (GFP-, mCherry+). Error bars indicate standard deviation. The numbers refer to the neuropil defined in **W, X.** 1 and 3: neuropils containing a *WT* (1) and a *Dll-/-* (2) MARCM clones, 2 and 4: contralateral neuropil with no clones.

Lin A, D and Z give rise on average to 79 (N=10), 57 (N=4) and 33 (N=13) astrocytes, respectively, but with a very high standard deviation, suggesting that each lineage generates a variable number of NG **(Figure S5)**. For each of the three lineages born in the T1 hemisegment, astrocyte processes mostly invade the ProNp (<u>P</u>rothoracic <u>N</u>euro<u>p</u>il) but, depending on the sample, can also send processes to the Accessory Mesothoracic Neuropil (AMesoNp, where wing MN dendrites innervate), the mesothoracic neuropil (MesoNp), and occasionally cross the midline to populate the contralateral neuropil. Although most T1-born astrocyte cell bodies remain in the ProNm (<u>P</u>rothoracic <u>N</u>euro<u>mere</u>), they can also end up in the Accessory Mesothoracic Neuromere (AMesoNm) and occasionally in the Mesothoracic Neuromere (MesoNm) **(Figure 8A-R, data not shown for Lin D, Supp. Table 3)**. Similarly, Lin A astrocytes born in a T2 and T3 hemisegment have the same ability to populate neighboring neuromeres **(Figure S6)**. These observations demonstrate that a neuropil can be invaded by astrocyte processes born in neighboring segments. For example, the AMesoNp is populated by glia born in either T1 or T2. Thus, unlike MNs, the number, final cell body position, and neuropil regions invaded by astrocyte processes born from any of the three NGBs are variable from animal to animal **(Figure 8, S6, Supp. Table 3)**.

Although we were unable to count the number of EG produced by Lin Z **(see Methods)**, Lin A gives rise to ~114 EG (N=4) with a low standard deviation (+/-6), suggesting that the number of EG produced by Lin A is similar in different animals. However, as with astrocytes, the neuropil region surrounded and invaded by Lin A EG is variable from animal to animal, and these glia can populate neighboring neuromeres and cross the midline **(Figure 8, S6 and Supp. Table 3)**.

In summary, each of the three NGBs produces a variable number of NG with different morphologies and final locations, but the same number of motor neurons, each with a stereotyped morphology.

### Final astrocyte number depends on inter-lineage competition

Although the number of NG produced by a single NGB (e.g. Lin A) can be highly variable, the total number of NG produced by all three NGBs is very constant from animal to animal **(Figure 1, S5)**. In addition, the non-stereotyped morphology and final positions of NG derived from single NGBs led us to hypothesize that NG may compete during development to ensure that each neuropil is fully wrapped and innervated. We tested this hypothesis for astrocytes, by generating WT or *Dll* mutant Lin A MARCM clones in a genetic background in which we could also count the total number of astrocytes. Because the number of astrocytes produced by *Dll* mutant Lin A is severely compromised **(Figure 7)**, we considered three possible outcomes **(Figure 8)**: (1) No compensation, in which WT (non-Lin A) astrocytes do not respond to the reduced number of astrocytes derived from Lin A; (2) Compensation where WT astrocytes respond to the lower number of Lin A astrocytes by increasing the number or extent of processes invading the neuropil; and (3) Compensation where WT astrocyte progenitors divide more to maintain the correct number of astrocytes **(Figure 8S-V)**.

To distinguish between these possibilities, we generated WT or *Dll* mutant Lin A MARCM clones expressing mCD8::GFP under the control of *repo-Gal4* and

*DVGLUT-Gal4*. In this genetic background, we labeled all astrocytes with a cytoplasmic QUAS-mCherry under the control of *alrm-QF*. As a result, astrocytes produced by Lin A are labeled by mCherry and mCD8::GFP while astrocytes produced by other lineages express only mCherry. Astrocytes were counted in a ProNm containing a single Lin A MARCM clone and in the contralateral neuromere that had no clones, which served as an internal control. Although *Dll* mutant Lin A clones (N=4) produced on average ~18 astrocytes (compared to ~80 for WT Lin A clones), the total number of astrocytes in these neuromeres was normal (~140) (**Figure 8S-Z**).

Altogether, these results support the idea that NG compete with each other during metamorphosis to ensure that they are generated in correct numbers.

## Discussion

### The logic of lineages producing multiple, functionally related cell types

Here, we show that three NGBs per thoracic larval hemisegment give rise to motor neurons, astrocytes and ensheathing glia. All three of these cell types are key components of each thoracic neuromere: MNs extend axons to innervate leg muscles and elaborate complex dendritic arbors inside the neuropil, ensheathing glia wrap the developing neuropil and incoming sensory axons, and astrocytes send processes inside the neuropil where they associate with synapses between MN dendrites, interneurons, and sensory neurons. Together, these cells develop in a coordinated manner to form a complex functional unit that comprises the neural circuitry used for many adult behaviors such as walking.

Glial cells are a major component of nervous systems, critical not only for the activity of neurons but also for their development (Freeman, 2015; Freeman and Rowitch, 2013). Neurons and glia are always associated with each other and exist in all bilateria, even in basal phyla such as flatworms (Hartline, 2011). As nervous system complexity increases, there is an increase in both neuronal diversity and in the diversity of glia types and morphologies (Paredes et al., 2016). Given this close relationship, it is striking that in the thorax of the adult fly all NG are derived from lineages that also (and *only*) give rise to leg MNs. In the absence of cell migration, being born from the same stem cell results in an anatomical proximity that may be important for subsequent steps in development. In particular, we propose that the shared lineages of the leg MNs, astrocytes, and EG facilitates the assembly of anatomically complex neuropils and neural circuits. This idea helps explain why lineage relationships appear to play less of a role in simpler nervous systems such as in *C. elegans*. Consistent with the idea that anatomical proximity of functionally related cell types is important for building complex nervous systems, in mammals, astrocytes and oligodendrocytes are born from the same progenitor domains as motor neurons and interneurons (Rowitch and Kriegstein, 2010; Ravanelli and Appel, 2015). Although it is currently not known if these cell types are derived from the same lineages in vertebrates, our results provide a striking precedent and compelling reasons that this may be the case. Alternatively, vertebrate nervous systems may have solved the neuropil assembly problem differently, by having distinct, temporally related, progenitors born in the same domain that give rise to either neurons or glia.

### Divergent mechanisms for generating MN and glia stereotypy

One of the most striking conclusions stemming from our findings is how cell lineages can use very different mechanisms to produce stereotyped outcomes. On the one hand, each MN is morphologically distinct, and born from a specific lineage and with a specific birth order, properties that stem from the unique TF code that they express. At the other extreme, based on the hundreds of TFs we have surveyed, individual astrocytes or EG appear to share the same TF code as all other astrocytes or EG, respectively, and can end up with different morphologies in multiple positions within or even in neighboring neuropils. Yet, despite this plasticity, the end result – ~280 NG evenly distributed throughout each thoracic leg neuropil – is highly stereotyped. Nevertheless, these two very different modes of achieving stereotypy – hardwired (for MNs) and plastic (for NG) – arise from the same stem cells. We also found that although the total number of NG in each thoracic neuropil is stereotyped (~140 astrocytes and ~140 EG), the individual contributions from the three NGB lineages can vary from animal to animal. Moreover, when one lineage is compromised the others can compensate to maintain the correct number of total NG. Although the mechanism regulating final NG number is currently unknown, we suggest that it is analogous to what has been referred to as neutral competition in mammalian and *Drosophila* gut homeostasis (de Navascues et al., 2012; Snippert et al., 2010). It is possible that this competition occurs via direct communication between NG. This idea is supported by the observation that individual astrocytes and EG respect each other’s territory, and thus exhibit tiling-like behavior. Alternatively, communication between MN dendrites (or other components of the thoracic neuropil) and NG may be critical for NG stereotypy. For example, astrocytes and EG may be able to sense and adapt to the scaffold generated by MN dendrites in order to fully invade and wrap each thoracic neuropil. Accordingly, NG stereotypy, both cell number and morphology, would be an indirect consequence of the mTF codes that generate the unique morphologies of MNs and likely other neurons.

What might be the reason for the existence of such different strategies to achieve stereotypy? One answer may be developmental robustness, which is especially challenging for establishing neural circuits. While MNs need to make precise connections (both in the CNS and in the legs), and therefore may require precise TF codes to achieve this precision, our results suggest that astrocytes and EG may have a more generic role in neuropil development. Nevertheless, NG are likely to be just as critical for neuropil development and function. However, instead of specifying each glia on a single cell basis, the system has evolved a different strategy to achieve robustness: a generic TF code, but the ability to communicate with each other to ensure that the entire neuropil is appropriately and evenly populated by the correct number of NG. There are several potential advantages to such a strategy. One is that as the number of MNs or size and complexity of nervous systems vary during evolution, the number of NG can readily adapt in response. Second, NG plasticity may be crucial during normal development to adjust to natural variations in neuronal morphology. For example, in the antennal lobe of Drosophila the morphology and connectivity of local interneurons can vary between animals, and may require NG to respond accordingly (Chou et al., 2010). Finally, neuronal numbers or morphologies may differ due to errors that occur during development or as a consequence of injury. Glial developmental plasticity may be essential to achieve robustness by readily adapting to these and perhaps other variations in nervous system size and morphology.

## Experimental procedure

### Fly Stocks

Unless otherwise noted, fly stocks were obtained from the Bloomington Stock Center: *alrm-Gal4-III*, *UAS-mCD8::GFP-III, UAS-mCD8::GFP-II UAS-H2B::RFP-III (Bernd Mayer et. Al, 2005), UASFB1.1-II, R56F03-Gal4*; UAS-GFP::CD8::HRP (Alexandre C et. Al 2014); *R56F03-Gal4, UASCD4-tdGFP recombinant ( Peco et al., 2016), UAS-mFlp5 (Enriquez et al. 2015); dll-Gal4, R31F10-Gal4, w hs-Flp^1.22^ FRT82B tub-Gal80 and FRT42D tub-Gal80 (Gary Struhl), UAS-mCD8::GFP, Mhc-RFP, VGlut-Gal4 (also called OK371-Gal4) (Mahr and Aberle, 2006), tub-QS, repo-Gal4,*

*UAS-mCD8::RFP, VGlut-QF* (this work)*, Dll-( Dll[SA1]* (Cohen and Jurgens, 1989)*, QUAS-mcherry-III, UAS-mcherry-III, alrm-QF-III, elav-Gal80-I, cha-Gal80* (Kitamoto, T. 2002).

### Immunostaining of L3 Larva

Inverted L3 larvae were fixed in 4% formaldehyde with PBS for 20 minutes. L3 larval CNS were dissected in PBS triton and incubated with primary antibodies for two days and secondary antibodies for one day at 4°C. Fresh PBT (PBS with 0.1% Triton

X-100 0,3%, 1% BSA) was used for the blocking step, incubation and washing steps: five times for 20 minutes at room temperature after fixation and after primary/secondary antibodies. L3 CNS mounted onto glass slides using Vectashield mounting medium (Vector Labs).

### Immunostaining of adult VNS

After removing the abdominal and head segments, the thoraces of the flies were opened and fixed in 4% formaldehyde with PBS for 20 minutes at room temperature and blocked in the blocking buffer for one hour. After dissection, adult VNS were incubated with primary antibodies for two days and secondary antibodies for one day at 4°C. Fresh PBT (PBS with 0.1% Triton x-100, 1% BSA) was used for the blocking step, incubation and washing steps: five times for 20 minutes at room temperature after fixation and after primary/secondary antibodies. VNSs were dissected and mounted onto glass slides using Vectashield mounting medium (Vector Labs).

### Primary and secondary antibodies

Unless otherwise noted primary antibodies were obtained from Hybridoma Bank: mouse anti-BRP, rat anti-Elav, mouse anti-Nrg (neuronal specific form), mouse anti-Pros, rat anti-NCad, guinea-pig anti-Dll (Estella, C et al., 2008), guinea-pig anti-Dpn (gift from Jim skeath). Secondary antibodies: Alexa 555, 647 (Thermofisher) and Alexa 405 (abcam).

### Time course of NG development

We expressed a membrane reporter (*UASmCD8::GFP*) with *Dll-Gal4* and dissected larva and pupae at different development time points **(Figure 3, video 3)**. In order to specifically visualize the immature leg neuropil, larval CNS were co-stained with anti-BRP (which labels mature neuropil) and anti-N-Cadherin (NCad, which labels mature and immature neuropil at high and low levels, respectively (Kurusu et al., 2012)) **(Figure 3A-E)**. *Dll-Gal4* is also expressed in adult leg sensory neurons, which allowed us to visualize where sensory axons first enter the CNS and was used as a reference point in these developing CNS. We also stained larval CNS for Dll and BRP or Repo and Dll in order to quantify the number of glia at each development stage **(Figure 3F-J and data not shown)**. Finally, we used *alrm-Gal4* and *R56FO3-Gal4* to drive the expression of mCD8::GFP to determine when EG and astrocytes morphologically differentiate.

### EM protocol

After removal of abdominal and head segments, the thorax of the flies were opened and fixed in 4% glutaraldehyde, washed 5 times quickly in PBS, incubate in PBS with 0,03% Triton X-100 for 30 mins and washed 5 times for 20 mins in PBS. To specifically recognize the EG in EM we expressed *UAS-GFP::mCD8::HRP (HRP: horseradish peroxidase)* to catalyze DAB (3,3′-Diaminobenzidine) and produce a precipitate dense to electrons. The expression of the HRP (Horseradish Peroxydase) were revealed overnight using the DAB substrate kit from Thermofisher.

With an EM microwave (Pelco model 3451 system with coldspot), specimens were post-fixed with 1% osmium tetroxide in 0.1M phosphate buffer (PB) for 2 x 40 sec (with each 40 second exposure in fresh osmium), and then washed 3 x 10 minutes in 0.1M PB. Specimens were dehydrated in ethanol grades of 50%, 70%, 95% (1 x 40 seconds for each grade), and 100% (2 x 40 seconds). Dehydrated tissue was then infiltrated with epoxy resin (Electron Microscopy Sciences Embed 812) and 100% ethanol (1:1 mixture) for 15 minutes, and then in 100% epoxy resin (2 x 15 minutes with fresh resin each time).

Specimens were mounted between 2 plastic slides with 100% epoxy resin and polymerized overnight at 60 degrees C. The next day the polymerized resin wafers were separated from the plastic slides and placed flat on a glass slide to choose specimens for thin sectioning. An entire specimen sample was then cut out of the wafer using a razor blade and carefully placed on its side in a Dykstra flat embedding mold with the protoracic neuromere (PN) end (region of interest) closest to the beveled tip in one of the cavities of the mold. Small slits are made the tip and floor of the silicon mold to secure the perpendicularly oriented tissue. This orientation is used so that thin sections will be cut transversely through the specimen starting at the PN. The mold is then filled with 100% epoxy resin and polymerized for 18-24 hours at 60 degrees C.

After polymerization, the block is removed from the cavity and trimmed down at the beveled tip in the shape of a 2mm square with the specimen near the center of the blockface. With an ultramicotome, 10 micron thick sections are cut and collected serially using a diamond Histo-knife (Diatome) and then mounted and coverslipped on a glass slide with immersion oil. The 10 micron sections are then observed with a light microscope and sections with the best areas of interest are photographed. The coverslip is carefully removed and the sections of interest are washed in 95% ethanol to remove any oil by passing them through the ethanol in 3 wells for 15 seconds in each well. Each 10 micron section is then carefully placed on a piece of lens tissue to dry and then remounted on a tiny drop of epoxy resin on a blank polymerized Beem capsule resin block (blockface should be flat and smooth) and polymerized overnight at 60 degrees C; a spring tension apparatus is used to keep the 10 micron section flat against a plastic slide. The polymerized blockface with the 10 micron section is trimmed down to size and 60 nanometer thin sections are cut with an ultramicrotome using a Diatome diamond knife. The ultrathin sections are collected onto formvar coated slot grids and then stained with uranyl acetate and lead citrate, then examined on a JEOL 1200EX electron microscope.

### Fly genetics

Crosses or stocks used to visualize NG (Figure 1, 2, 3 S1, S3):

*UAS-mCD8::GFP-II, alrm-Gal4-III*

*alrm-Gal4, UAS-FB1.1-III* cross with *UAS-mFlp5-II*

*alrm-Gal4, UAS-H2B:: RFP-III*

*UAS-mCD8::GFP-II, R56F03-Gal4-III*

*R56F03, UAS-FB1.1-III* cross with *UAS-mFlp5-II*

*R56F03, UAS-H2B:: RFP-III*

*R56F03-Gal4-III cross with UAS-GFP::CD8::HRP*

R56F03-Gal4, UAS-nGFP-III

R56F03-Gal4, UAS-nGFP-III cross *with alrm-Gal4*

*Dll-Gal4, UAS-mCD8::GFP-II*

Lineage tracing systems (Figure 2, S2):

*Elav-Gal80; Dll-Gal4, UAS-Flp1/cyo; cha-Gal80/MKRS cross with act>cd2>Gal4, UAS-nGFP-III.*

*Note:* since Dll-Gal4 is expressed in sensory neurons we used *cha-Gal80 and elav-Gal80* to inhibit Gal4 in these neurons.

*Elav-Gal80; dll-Gal4, UAS-Flp1/Cyo; cha-gal80/MKRS cross with UAS-mFlp5; UAS-FB1.1//; act>CD2>Gal4*

*R31F10-Gal4 cross with UAS-mFlp5-X, UAS-Flybow-I.I*

MARCM (Figure 4, 5):

*Y, w, hs-Flp^1.22^; FRT42 tub-gal80//; repo-Gal4, UAS-mCD8::GFP/MKRS* cross with

*y, w, hs-Flp^1.22^; VGlut-Gal4, UAS-mCD8::GFP, Mhc-RFP FRT42D//; repo-Gal4, UAS-mCD8::GFP/Tm6b or*

*y, w, hs-Flp^1.22^; VGlut-Gal4, UAS-mCD8::GFP, Mhc-RFP FRT42D Dll-//; repo-Gal4, UAS-mCD8:: GFP/Tm6b*

*y, w, hs-Flp^1.22^*, *tub-Gal4; sp/cyo; repo-gal4, FRT82B tub-Gal80 cross with*

*y, w, hs-Flp^1.22^*, UAS-mCD8::GFP//*; FRT82B//*

*y, w, hs-Flp^1.22^; FRT42D tub-Gal80 //; alrm-QF, QUAS-mcherry/MKRS cross with*

*y, w, hs-Flp^1.22^; VGlut-Gal4, UAS-mCD8::GFP, Mhc-RFP FRT42D Dll-//; repo-Gal4, UAS-mCD8::GFP/Tm6b* or

*y, w, hs-Flp^1.22^; VGlut-Gal4, UAS-mCD8::GFP, Mhc-RFP FRT42D //; repo-Gal4, UAS-mCD8:: GFP/Tm6b*

*cis^2^* MARCM (Figure 4, 6):

*y, w, hs-Flp^1.22^; FRT42 tub-Gal80, tub-QS/cyo; repo-Gal4, UAS-mCD8::RFPTM6b or*

*y, w, hs-Flp^1.22^; FRT42 tub-Gal80, tub-QS/cyo; R56F03-Gal4, UAS-mCD8::RFP/TM6b or*

*y, w, hs-Flp^1.22^; FRT42 tub-Gal80, tub-QS/cyo; alrm-Gal4, UAS-mCherry/TM6b or*

*y, w, hs-Flp^1.22^; FRT42 tub-Gal80, tub-QS/cyo; repo-Gal4, UAS-H2B::RFP, UASmCD8::GFP/TM6b or*

*y, w, hs-Flp^1.22^; FRT42 tub-Gal80, tub-QS/cyo; R56F03-Gal4, UAS-H2B::RFP/tm6b* or

*y, w, hs-Flp^1.22^; FRT42 tub-Gal80, tub-QS/cyo; alrm-Gal4, UAS-H2B::RFP/tm6b cross with*

*y, w, hs-Flp^1.22^; QUAS-mCD8: GFP FRT42D //; VGlut-QF/MKRS or*

*y, w, hs-Flp^1.22^; QUAS-mCD8:GFP FRT42D Dll-//; VGlut-QF/MKRS*

Note: *cis^2^* MARCM and MARCM clones where generated by heat shocks of first instar larva.

#### Cloning (Figures 4, 6, 7)

A SpeI-VGlut-Spe1 fragment containing the 5.9 kb sequence upstream from the *VGlut* translation start site was taken from a pCR™8/GW/TOPO (Enriquez et al.) vector and clone in a *pattb-QF-hsp70* vector (addgene). The resulting *pattb-VGlut-QF-hsp70* construct was inserted in position 86f on chromosome III by injection into embryos carrying *attP-86f* landing site.

#### Leg preparation (Figure 4)

Legs were dissected and fixed overnight at 4°C, washed five times for 20 minutes at room temperature in PBS withTriton X-100 (0.3%) and mounted onto glass slides using Vectashield mounting medium (Vector Labs).

#### Microscopy and 2D Imaging

Multiple 1-μm-thick sections in the z axis (dorsoventral for L3 CNS or adult VNS and mediolateral for adult legs) were imaged with a Leica TCS SP5 II or a zeiss LSM 780 confocal microscope. Binary images for z stack images were generated using NIH Image J and Photoshop (Adobe Systems).

#### 3D Leg analysis (Figure 4)

Z-stacks were generated using NIH Image J. Specific cuticle background was generated using the Argon laser 488 and by placing the confocal detector out of the range of the GFP emission wavelength. This cuticle background of the leg was used to remove the nonspecific signal from the GFP channel using ImageJ. Z-stacks were used to generate 3D reconstructions of legs with Amira 3D software. The color codes used were Volrengreen to visualize mCD8::GFP, color used for cuticles was grey.

#### 3D images of adult VNS and larval CNS

Ventral to dorsal Z-stacks were generated using NIH Image J. and were used to generate 3D reconstructions using the Volren or volume rendering in Amira 3D software. The color codes used were constant color in correlation with the fluorescent proteins or the secondary Alexa used.

## Acknowledgements

We thank members of the Mann lab for comments and suggestions, and Oliver Hobert, Emilie Peco, and Alain Vincent for comments on the manuscript. This work was supported by an NIH grant NS070644 to R.S.M. and funding from the ALS Association (#256) to J.E.

